# Enhanced functional divergence of duplicate genes several million years after gene duplication in the *Arabidopsis* lineage

**DOI:** 10.1101/047639

**Authors:** Kousuke Hanada, Ayumi Tezuka, Masafumi Nozawa, Yutaka Suzuki, Sumio Sugano, Atsushi J. Nagano, Motomi Ito, Shi-Ichi Morinaga

**Affiliations:** Frontier Research Academy for Young Researchers, Department of Bioscience and Bioinformatics, Kyusyu Institute of Technology, Iizuka, Fukuoka 820-8502, Japan; College of Bioresource Sciences, Nihon University, 1866 Kameino, Fujisawa, Kanagawa 252-0880, Japan; Graduate School of Frontier Science, The University of Tokyo, Kashiwa, Chiba 277-8562, Japan; Center of Ecological Research, Kyoto University, 2-509-3, Hirano, Otsu, Shiga 520-2113, Japan; Graduate School of Arts and Sciences, The University of Tokyo, Komaba 3-8-1, Tokyo 153-8902, Japan; RIKEN Center for Sustainable Resource Science, RIKEN, Yokohama, Kanagawa 230-0045, Japan; CREST, Japan Science and Technology Agency, Kawaguchi, Saitama 332-0012, Japan; Center for Information Biology, National Institute of Genetics, 1111 Yata, Mishima,Shizuoka 411-8540, Japan; Department of Genetics, SOKENDAI, 1111 Yata, Mishima, Shizuoka 411-8540, Japan

**Author notes:** **Kousuke Hanada**, Frontier Research Academy for Young Researchers, Kyushu Institute of Technology, 680-4 Kawazu, Iizuka, Fukuoka 820-8502, Japan; **Shin-Ichi Morinaga**, College of Bioresource Sciences, Nihon University, 1866 Kameino, Fujisawa, Kanagawa 252-0880, Japan.

**Keywords:** *Arabidopsis*, Gene duplication, Positive selection

## Abstract

Lineage-specifically duplicated genes likely contribute to the phenotypic divergence in closely related species. However, neither the frequency of duplication events nor the degree of selective pressures immediately after gene duplication is clear in the speciation process. Plants have substantially higher gene duplication rates than most other eukaryotes. Here, using Illumina short reads from *Arabidopsis halleri*, which has highly qualified plant genomes in close species (*Brassica rapa, A. thaliana* and *A. lyrata*), we succeeded in generating orthologous gene groups among *B. rapa, A. thaliana, A. lyrata* and *A. halleri*. The frequency of duplication events in the *Arabidopsis* lineage was approximately 10 times higher than the frequency inferred by comparative genomics of *Arabidopsis*, poplar, rice and moss. Of the currently retained genes in *A. halleri*, 11–24% had undergone gene duplication in the *Arabidopsis* lineage. To examine the degree of selective pressure for duplicated genes, we calculated the ratios of nonsynonymous to synonymous substitution rates (*K*_A_/*K*_S_) in the *A. halleri-lyrata* and *A. halleri* lineages. Using a maximum-likelihood framework, we examined positive (*K*_A_/*K*_S_ > 1) and purifying selection (*K*_A_/*K*_S_ < 1) at a significant level (P < 0.01). Duplicate genes tended to have a higher proportion of positive selection compared with non-duplicated genes. More interestingly, we found that functional divergence of duplicated genes was accelerated several million years after gene duplication at a higher proportion than immediately after gene duplication.

## Introduction

Plant genomes have experienced more gene duplication events than most other eukaryotes (Lockton and Gaut 2005). These duplications are largely classified into whole genome duplications (WGDs) and small-scale duplications (SSDs). WGD events are concentrated in the Cretaceous-Paleogene extinction period (Vanneste et al. 2014). SSDs such as retroduplication and tandem duplication have occurred continuously at a high rate during plant evolution, indicating that SSDs may be a key factor for plant speciation or phenotypic differences in close relatives (Fortna et al. 2004; Rostoks et al. 2005; Rizzon et al. 2006; Clark et al. 2007; Hanada et al. 2008; Hanada et al. 2009c). SSDs tend to be lineage-specific in land plants (Hanada et al. 2008). There is a clear functional bias between WGD and SSD in plants (Hanada et al. 2008). SSDs tend to be associated with stress responses (Rizzon et al. 2006; Hanada et al. 2008), and are likely important for adaptive evolution to rapidly changing environments in close relatives.

Lineage-specifically duplicated genes, as a typical example of SSDs, have been identified between *Arabidopsis thaliana* and *A. lyrata*, whose complete genome sequences have been determined (Arabidopsis_Genome_Initiative. 2000; Hu et al. 2011). One-hundred and thirty-seven genes duplicated after the divergence of *A. thaliana* and *A. lyrata* 10–11 million years ago (MYA) (Moghe et al. 2014) were identified in the *A. thaliana* lineage through careful analysis (Wang et al. 2013). These lineage-specifically duplicated genes tend to have more divergent expressional profiles. Such duplicated genes may have contributed to the functional divergence between *A. thaliana* and *A. lyrata*. Although recent studies have succeeded in constructing many qualified genomes using mate-paired and paired-end Illumina short reads, great reading depth is necessary to generate a qualified genome.

*Arabidopsis* has two qualified genomes in *A. thaliana* and *A. lyrata. A. halleri* is a close relative of both, but a qualified genome for *A. halleri* is unavailable. Using Illumina paired-end short reads from *A. halleri* and the two qualified genomes, we inferred orthologous genes against the *A. thaliana* and *A. lyrata* genes without generating an *A. halleri* qualified genome. As an outgroup species for the three *Arabidopsis* species, we used *Brassica rapa*, which also has a qualified genome sequence. We generated three sets of orthologous gene groups (OGGs) among *B. rapa, A. thaliana, A. lyrata* and *A. halleri* (**Fig. 1**). The divergence time between *A. lyrata* and *A. halleri* is supposed to be approximately 2 MYA (Beilstein et al. 2010), and the divergence time between either *A. halleri* or *A. lyrata* and *A. thaliana* is approximately 10–11 MYA (Moghe et al. 2014). Here, the *A. halleri-lyrata* lineage was defined as the common ancestor lineage between *A. halleri* and *A. lyrata* after the divergence of *A. thaliana* (**Fig. 2**). The *A. halleri* lineage was defined as the lineage after the divergence of *A. lyrata* (**Fig. 2**). There are phenotypic variations among the three species such as self-compatibility and heavy-metal tolerance. *A. thaliana* has self-compatibility, but the others are self-incompatible (Verbruggen et al. 2009; Kramer 2010; Koenig and Weigel 2015). Therefore, *A. halleri* and *A. lyrata* have large petals to attract pollinator insects, and the anthers are separated from the stigma to avoid autopollination (Shimizu and Purugganan 2005). *A. halleri* is known as a Zn/Cd hyper-accumulator (Verbruggen et al. 2009; Kramer 2010) and some wild populations of *A. lyrata* are tolerant of serpentine soils, which are characterized by a high heavy-metal content (Turner et al. 2010), while *A. thaliana* is not.

**Figure 1.**
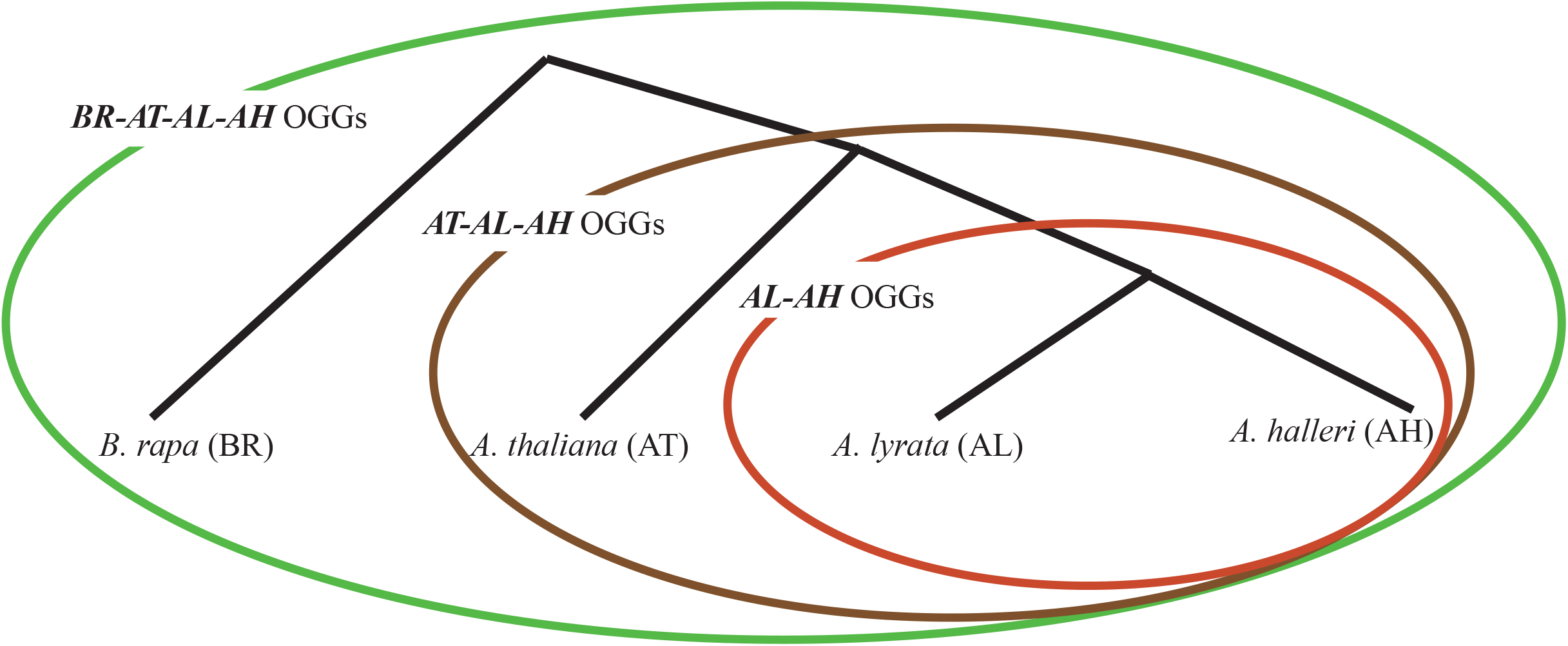
**Three sets of orthologous gene groups among *B. rapa, A. thaliana, A. lyrata* and *A. halleri*** Orthologous gene groups between *A. lyrata* and *A. halleri* were defined as *AL-AH* OGGs (red solid line). There were 28,036 *AL-AH* OGGs as shown in Supplemental Fig. S1. Orthologous gene groups among *A. thaliana, A. lyrata* and *A. halleri* were defined as *AT-AL-AH* OGGs (brown solid line). There were 23,086 *AT-AL-AH* OGGs as shown in Supplemental Fig. S2. Orthologous gene groups among *B. rapa, A. thaliana, A. lyrata* and *A. halleri* were defined as *BR-AT-AL-AH* OGGs (green solid line). There were 17,641 *BR-AT-AL-AH* OGGs as shown in Supplemental Fig. S3.

**Figure 2.**
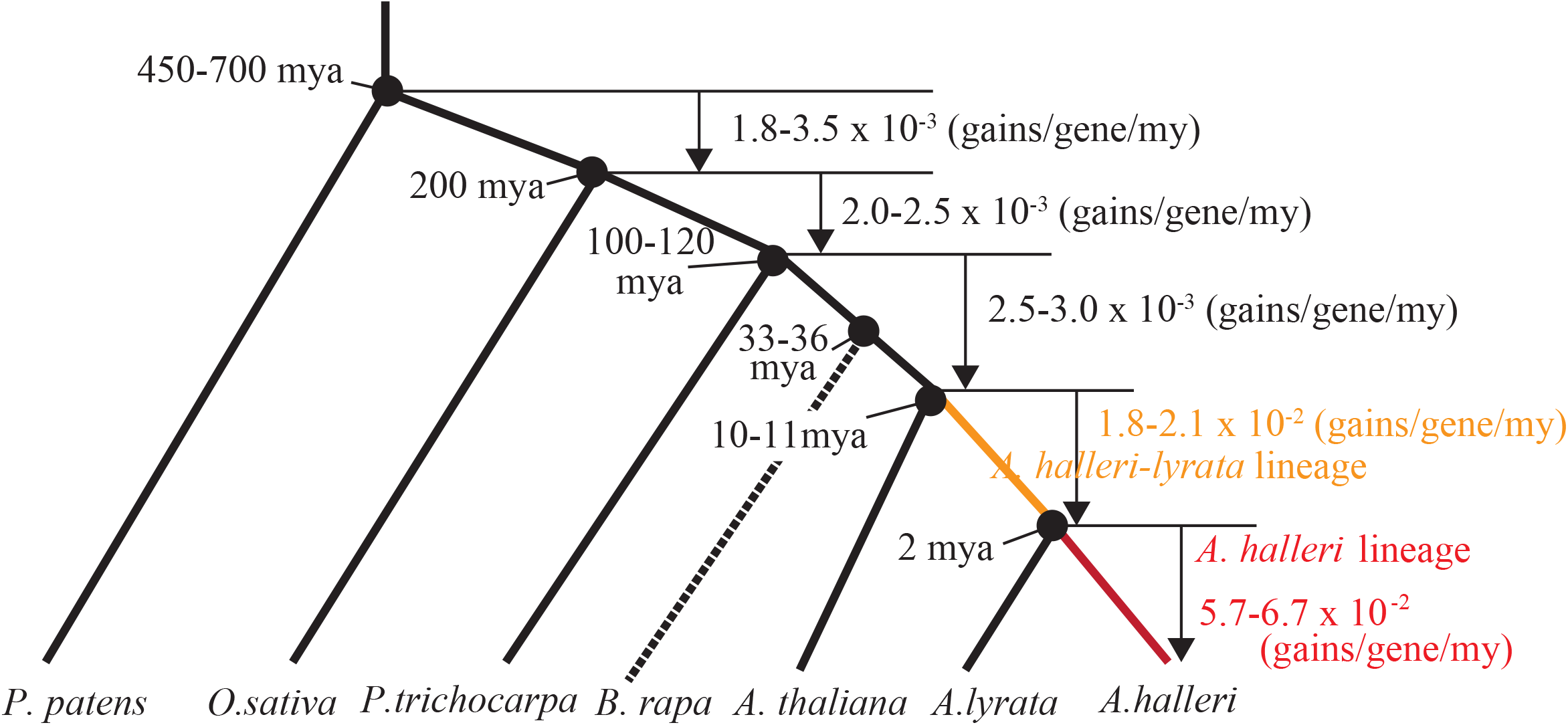
**Gain rates through gene duplication in land plants** The divergence times among mosses (*Physcomitrella patens*), rice (*Oryza sativa*), poplar (*Populus trichocarpa*), *B. rapa, A. thaliana, A. lyrata* and *A. halleri* were taken from previous reports (Wolfe et al. 1989; Tuskan et al. 2006; Rensing et al. 2008; Beilstein et al. 2010; Heckman and Kauffmann 2011; Moghe et al. 2014). Gene gains through gene duplication were estimated for each branch. The gene gains in two branches were estimated in the present study. One was the *A. halleri-lyrata* lineage (orange line), which represents the evolutionary process of the *A. halleri* lineage between the divergence of *A. thaliana* and the divergence of *A. lyrata*. The other was the *A. halleri* lineage (red line), which represents the evolutionary process of the *A. halleri* lineage after the divergence of *A. lyrata*. Gene gains were divided by ancestral gene numbers and branch lengths corresponding to times (million years). The rate of gene gain was defined as the number of genes gained through gene duplication per gene per million years.

Duplicated genes have contributed to functional divergence during plant evolution (Hanada et al. 2009b). The gene dosage hypothesis proposes that duplicate genes can be fixed for increased gene dosage, keeping redundant functions among duplicated genes. In *A. halleri*, the expression of HEAVY METAL ATPASE 4 (HMA4) has been enhanced by cis-regulatory mutations and increased gene copy number for metal hyperaccumulation (Hanikenne et al. 2008). This is an example of increased gene dosage by gene duplication. The degree of functional divergence or constraint can be inferred by selection pressures based on the nonsynonymous substitution rate (*K*_A_) versus the synonymous substitution rate (*K*_S_). Genes undergoing positive selection (*K*_A_/*K*_S_ > 1) and purifying selection (*K*_A_/*K*_S_ < 1) are associated with functional divergence and constraint, respectively. Consequently, duplicated genes with purifying selection tend to be associated with increased gene dosage.

To understand the contribution of duplicated genes to the speciation process in *Arabidopsis* within the last 10 million years (MY), we first examined the frequency of gene duplications in the *A. halleri-lyrata* and *A. halleri* lineages. We then examined the relationship between gene duplication and selection pressures based on the *K_A_/K_S_* ratio. Finally, we examined the overrepresented functional categories of genes under positive and purifying selection.

## Results and Discussion

### Gene assembly of A. halleri

We generated *A. halleri* contigs from a total of 44.5 Gb Illumina sequencing reads using several assembly software tools. The *A. halleri* contigs generated by the ABySS software had the highest mapping rate (72% = 23,681/32,670) to 32,670 annotated *A. lyrata* genes with more than 80% similarity and 80% coverage. The best-matched region of an *A. halleri* contig was defined as the orthologous *A. halleri* gene sequence to an *A. lyrata* gene. However, an *A. lyrata* gene sequence was frequently mapped to different *A. halleri* contigs depending on the region of the *A. lyrata* gene because the whole sequence of each *A. halleri* gene was split into several contigs. To further identify *A. halleri* genes orthologous to *A. lyrata*, we re-assembled the *A. halleri* contigs based on the mapping information of *A. lyrata* genes. Through this, we obtained an additional 4,355 *A. halleri* gene sequences with more than 80% similarity and 80% coverage against *A. lyrata* genes. Given that all identified *A. halleri* genes were paired with *A. lyrata* genes, a total of 28,036 pairs were defined as *A. lyrata–A. halleri* (*AL*-*AH)* orthologous gene groups. The analysis procedures and findings are summarized in **Supplemental Figure S1**. The constructed *A. halleri* gene sequences with annotated *A. lyrata* gene IDs are shown in **Supplemental Table S1**.

### Duplication frequency in the A. halleri-lyrata lineage

To examine the frequency of duplication events in the *A. halleri-lyrata* lineage (**Fig. 2**), we searched for an orthologous *A. thaliana* gene for each *AL*-*AH* OGG by BLASTP searches between *AL*-*AH* OGG protein sequences and annotated *A. thaliana* protein sequences (Boratyn et al. 2013). When both the *A. lyrata* and *A. halleri* genes in an *AL*-*AH* OGG had the best hit to the same *A. thaliana* gene, the *A. thaliana* gene was defined as an orthologous gene. Of 28,036 *AL*-*AH* OGGs, 88% (24,784/28,036) had orthologous candidate genes in *A. thaliana* (**Supplemental Table S1**). To examine whether the candidate orthologous *A. thaliana* genes belonged to the *AL*-*AH* OGGs, we calculated synonymous substitution rates among the *A. thaliana, A. lyrata* and *A. halleri* genes. Of 24,784 potential OGGs among *A. thaliana, A. lyrata* and *A. halleri*, 23,086 (23,086/24,956 = 92.5%) were defined as *A. thaliana*-*A. lyrata*-*A. halleri* (*AT*-*AL*-*AH)* OGGs in which *A. thaliana* genes were outgroups of the *A. lyrata* and *A. halleri* genes. The analysis procedures and findings are summarized in **Supplemental Figure S2**.

In the 23,086 *AT*-*AL*-*AH* OGGs, 17,566 and 2,252 *A. thaliana* genes were uniquely and multiply assigned to *AT*-*AL*-*AH* OGGs, respectively. That is, the 23,086 *AT*-*AL*-*AH* OGGs were derived from 19,818 genes, which represent the number of genes in the common ancestor of *A. thaliana, A. lyrata* and *A. halleri*. Thus, 3,268 (23,086 - 19,818) gains through gene duplication were supposed to have occurred in the *A. halleri-lyrata* lineage. The gain rate (total gene duplication gains during a given time period divided by the estimated duration per gene) was inferred to be 1.8–2.1 × 10^-2^ (3,268 gains/19,818 genes/8–9 MY) in the *A. halleri-lyrata* lineage (**Fig. 2**).

In our previous study, we inferred the gain rate of gene duplication in the lineage leading to *Arabidopsis* after the divergence of mosses (Hanada et al. 2008). Using the same method, we re-estimated the gain rates in three times periods after the divergence of mosses (*Physcomitrella patens*), rice (*Oryza sativa*) and poplar (*Populus trichocarpa*). The gain rates were 1.8–3.0 × 10^-3^ in the three branches, approximately 10 times lower than in the *A. halleri-lyrata* lineage (**Fig. 2**). One explanation for this gain rate difference is that, immediately after speciation, the duplication rate was substantially higher. Another explanation is that even though many genes were fixed and retained, a large number of them did not survive in the long run. The latter explanation is consistent with the gradual decay of paralog synonymous substitution rates observed in several eukaryotes over time (Lynch and Conery 2000).

### Duplication frequency in the A. halleri lineage

Copy numbers of the *A. halleri* genes were unclear from *AT-AL-AH* OGGs. The absolute copy number of an *A. halleri* gene could be experimentally inferred by the absolute DNA concentration of the gene by droplet digital PCR (ddPCR) (Heredia et al. 2013). First, to examine the relationship between copy number and DNA concentration, we focused on a known three-copy gene, HMA4 (Hanikenne et al. 2008), and 11 singleton genes that share a single copy across four species from a broad range of plants (Duarte et al. 2010). The concentration of HMA4 was 627.5 ± 40.39 and 626.75 ± 39.38 copies/μl (four replicates for each of the two primer pairs, average of each primer pair ± 95% CI), whereas the concentration of the 11 singleton genes was 194.43 ± 5.50 copies/μl (four replicates for each of the 11 primer pairs, average of all 11 primer pairs ± 95% CI). The average DNA concentration was approximately 3.2 times higher for the three-copy gene compared with the single-copy genes (**Fig. 3A**), indicating that the copy number corresponded to the DNA concentration inferred by ddPCR.

**Figure 3.**
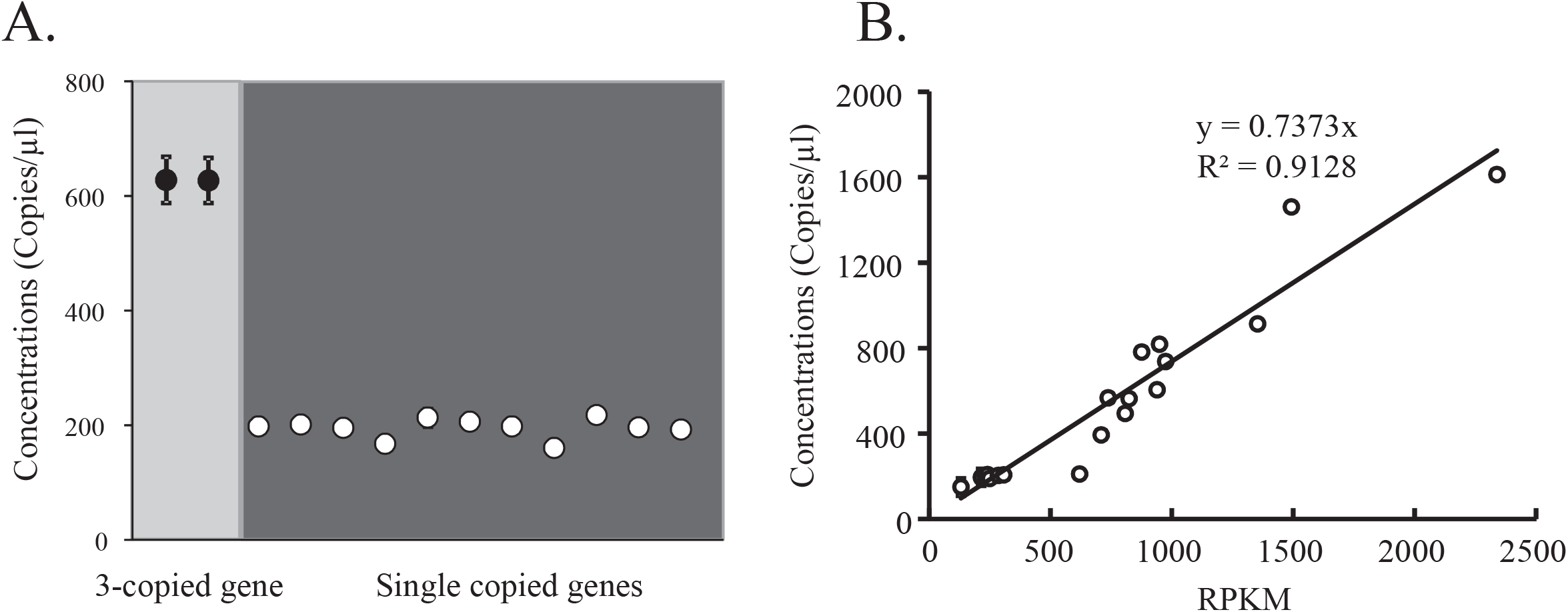
**Relationship between the Illumina sequencing read depth and the copy number inferred by ddPCR** (A) The Y-axis represents copy numbers per μl inferred by ddPCR. Black circles (grey background) and open circles (black background) indicate three-copy genes and single-copy genes, respectively. All points and error bars represent averages of four replicates and 95% confidence intervals. (B) Each dot represents an *A. halleri* gene. The X-axis represents the Illumina sequencing read depth, which is the number of reads per 1 Kbp per 1 million reads. The Y-axis represents copy numbers per μl, which were inferred by ddPCR. The regression line was calculated with the simple formula Y = αX; α was inferred by the least squares method.

The copy numbers of the *A. halleri* genes could also be inferred by the reading depth of the Illumina short reads. The reading depth was defined as the Reads Per Kilobase of exon per Million mapped reads (RPKM). To examine whether RPKM values corresponded with copy numbers, we focused on 27 *A. halleri* genes that had a wide variety of RPKMs (**see Materials and Methods**). For these genes, the copy number estimated by digital droplet PCR (ddPCR) was compared with the RPKM. Consequently, we found that RPKM was significantly correlated with concentration (**Fig. 3B, *R*^2^ = 0.91**). This result indicated that RPKM values could be used an index of copy numbers for *A. halleri* genes. Because of the 5% false positive rate when calling duplicated genes, we focused on 959 single-copy genes among a broad range of plant species (Duarte et al. 2010). The top 5% (298.38) of RPKMs of the 959 genes was defined as the threshold for duplicated genes. That is, *A. halleri* genes with an RPKM < 298.38 were defined as non-duplicated genes. *A. halleri* genes with RPKM values from 298.38 to 596.76 (298.38 × 2) were defined as two-copy genes, and *A. halleri* genes with RPKM values of 596.76 to 895.14 (298.38 × 3) RPKM were defined as three-copy genes. Following this rule, we identified 2,558 multiply duplicated *A. halleri* genes, and 3,741 gain events through gene duplications in the *A. halleri* lineage. The gain rate (total gains from gene duplication during a given time period divided by the estimated duration per gene) was 6.7 × 10^-2^ (3,741 gains/23,086 genes/2 MY) (Fig. 2). However, the gain events may have been over-estimated because duplication events occurring in the *A. halleri-lyrata* lineage caused an increase of RPKMs. Therefore, excluding *A. halleri* genes that were duplicated in the *A. halleri-lyrata* lineage, we counted the number of gain events in the *A. halleri* lineage. From 17,566 non-duplicated *A. halleri* genes in the *A. halleri-lyrata* lineage, the number of gain events was inferred to be 2,017 in the *A. halleri* lineage. The re-estimated gain rate was 5.7 × 10^-2^ (2,017 gains/17,566 genes/2 MY). The gain rate of duplicate genes in the *A. halleri* lineage was approximately two to three times higher than in the *A. halleri-lyrata* lineage. This result supports the rapid decay of duplicated genes over time (Lynch and Conery 2000).

### Selective pressures in the A. halleri-lyrata lineage

We found that 11–24% of OGGs had undergone gene duplications in the *Arabidopsis* lineage at high gain rates (1.8–6.7 × 10^-2^ gains/gene/MY), which were 10–30 times higher than the rates inferred by comparative genomics among *Arabidopsis*, poplar, rice and mosses (**Fig. 2**). Threfore, we were interested in investigating the functional divergence of duplicated genes in *Arabidopsis*. To infer the functional divergence of duplicated genes in the *A. halleri-lyrata* lineage, we tried to infer the ancestral sequences of the most recent common ancestor of *A. thaliana, A. lyrata* and *A. halleri*. To define the node of the most recent common ancestor in each *AT*-*AL*-*AH* OGG, we used an orthologous *Brassica rapa* gene as an outgroup sequence for 23,086 *AT*-*AL*-*AH* OGGs. We performed BLASTP searches of the *AT*-*AL*-*AH* OGG protein sequences against all *B. rapa* protein sequences (Boratyn et al. 2013). When the best-hit *B. rapa* gene was the same for the *A. thaliana, A. lyrata* and *A. halleri* genes, the *B. rapa* gene was considered a candidate orthologous gene to the *AT*-*AL*-*AH* OGG. After generating a phylogenetic tree for each OGG including *A. thaliana, A. lyrata, A. halleri* and *B. rapa* genes, we found 17,641 OGGs among *B. rapa, A. thaliana, A. lyrata* and *A. halleri* that were consistent with the topology of the species tree (**Supplemental Table S1**). These OGGs were defined as *BR-AT-AL-AH* OGGs. The analysis procedures and findings are summarized in **Supplemental Figure S3**.

Based on the phylogenetic tree in each *BR-AT-AL-AH* OGG, we inferred the ancestor sequences of all nodes using codeml in the PAML package (Yang 2007), and calculated *K*_A_ and *K*_S_ in the *A. halleri-lyrata* lineage. Among the 17,641 OGGs, we found 939, 8,279, and 8,423 genes with positive selection (*K*_A_/*K*_S_ > 1), neutral evolution and purifying selection (*K*_A_/*K*_S_ < 1), respectively, by the maximum likelihood approach. To address whether the duplicated genes contributed to functional divergence in the *A. halleri-lyrata* lineage, the proportion of positive selection, neutral evolution or purifying selection in the duplicated genes was compared with the proportion of positive selection, neutral evolution or purifying selection in non-duplicated genes. In this test, the null model was the ratio of duplicated genes to non-duplicated genes in the *A. halleri-lyrata* lineage. The proportion of either positive selection or neutral evolution in the duplicated genes was significantly higher than the proportion of either positive selection or neutral evolution in the non-duplicated genes (*P*-value = 2.2 × 10^-16^ for positive selection, *P*-value = 0.003 for neutral evolution by χ^2^ test, Fig. 4A). The proportion of purifying selection in the non-duplicated genes was significantly higher than the proportion of purifying selection in the duplicated genes in the *A. halleri-lyrata* lineage (*P*-value = 0.003 by χ^2^ test, Fig. 4A). These results indicate that gene duplication induced functional divergence in the *A. halleri-lyrata* lineage.

**Figure 4.**
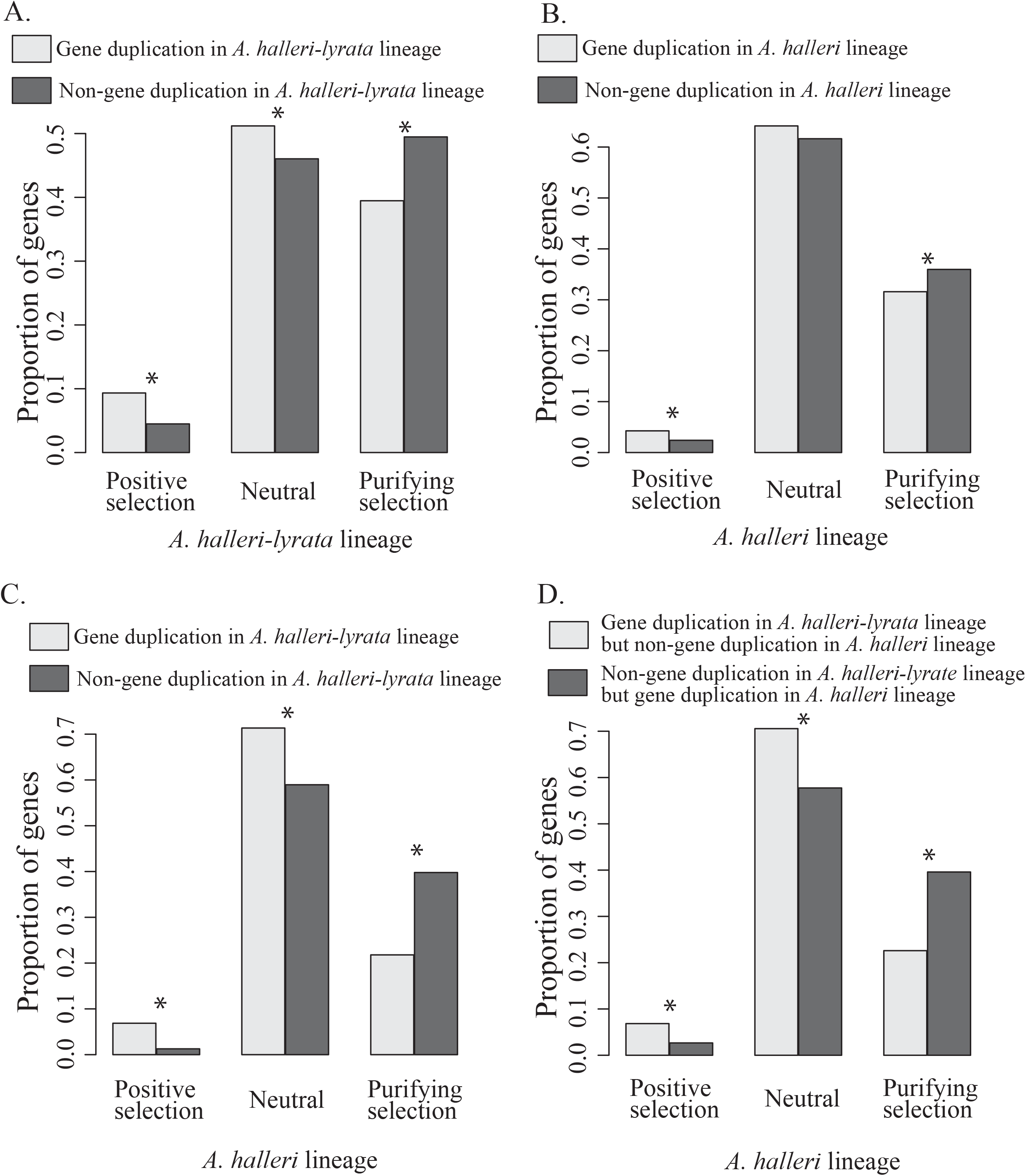
**Gene duplication and selective pressure in the *A. halleri-lyrata* and *A. halleri* lineages** Genes were classified as being under positive selection (K_A_/K_S_>1), neutral evolution (K_A_/K_S_=1) or purifying selection (K_A_/K_S_<1) in the *A. halleri-lyrata* and *A. halleri* lineages. Asterisks (*) represent significant differences by the chi-square test (*P*<0.01). (A) The relationship between gene duplication and selective pressure in the *A. lyrata-halleri* lineage. (B) The relationship between gene duplication and selective pressure in the *A. halleri* lineage. (C) The relationship between gene duplication in the *A. lyrata-halleri* lineage and selective pressure in the *A. halleri* lineage. (D) Comparison of selective pressure in the *A. halleri* lineage between gene duplication in only the *A. lyrata-halleri* lineage and gene duplication in only the *A. halleri* lineage.

### Selective pressure in the A. halleri lineage

To infer selective pressure in the *A. halleri* lineage, we generated ancestral sequences of the *A. lyrata* and *A. halleri* genes using *A. thaliana* genes as outgroups. When an *A. halleri* gene did not have any sequence variation in the Illumina short reads, *K*_S_ and *K*_A_ were simply calculated by comparing the ancestral sequence to the representative *A. halleri* gene sequence. When sequence variation for a representative *A. halleri* gene sequence was identified from the Illumina short reads, *K*_S_ and *K*_A_ were separately calculated for the varied sequences (see Materials and Methods, **Fig. S4**). Among the 23,086 *AL*-*AH* OGGs, we found 602, 14,293, and 8,191 genes with positive selection (*K*_A_/*K*_S_ > 1), neutral evolution and purifying selection (*K*_A_/*K*_S_ < 1), respectively, in the *A. halleri* lineage by the maximum likelihood approach. To examine the relationship between the effect of gene duplication and selective pressures in the *A. halleri* lineage, the proportion of positive selection, neutral evolution or purifying selection in the duplicated genes was compared with the proportion of positive selection, neutral evolution or purifying selection in the non-duplicated genes. In this test, the null model was the ratio of duplicated genes to non-duplicated genes in the *A. halleri* lineage. As observed in the *A. halleri-lyrata* lineage, the proportion of positive selection in the duplicated genes was significantly higher than the proportion of positive selection in the non-duplicated genes (*P*-value = 1.0 × 10^-7^ by χ^2^ test, Fig. 4B), while the proportion of purifying selection in the non-duplicated genes was significantly higher than the proportion of purifying selection in the duplicated genes (*P*-value = 0.002 by χ^2^ test, Fig. 4B), indicating that gene duplication induced functional divergence in the *A. halleri* lineage as well.

Functional divergence may not occur immediately after gene duplication. To determine whether gene duplication in the *A. halleri-lyrata* lineage affected selective pressures in the *A. halleri* lineage, we classified the 23,086 AL-AH OGGs into 5,520 OGGs with gene duplications and 17,566 OGGs without any gene duplications, focusing on the *A. halleri-lyrata* lineage. The proportion of positive selection, neutral evolution or purifying selection in the duplicated genes in the *A. halleri* lineage was compared with the proportion of positive selection, neutral evolution or purifying selection in the non-duplicated genes. In this test, the null model was the ratio of duplicated genes to non-duplicated genes in the *A. halleri-lyrata* lineage. The proportion of either positive selection or neutral evolution in the duplicated genes was significantly higher than the proportion of either positive selection or neutral evolution in the non-duplicated genes (*P*-value = 2.2 × 10^-16^ for positive selection and 3.9 × 10^-15^ for neutral evolution by χ^2^ test, Fig. 4C). The proportion of purifying selection in the non-duplicated genes was significantly higher than the proportion of purifying selection in the duplicated genes (*P*-value = 2.2 × 10^-16^ by χ^2^ test, Fig. 4C). These results indicate that gene duplication in the *A. halleri-lyrata* lineage contributed to functional divergence in the *A. halleri* lineage.

To determine whether gene duplications in the *A. halleri-lyrata* or *A. halleri* lineage contributed to functional divergence in the *A. halleri* lineage, we focused on two categories of OGGs—genes not duplicated in the *A. halleri-lyrata* lineage but duplicated in the *A. halleri* lineage (1,614 OGGs), and genes duplicated in the *A. halleri-lyrata* lineage but not duplicated in the *A. halleri* lineage (4,576 OGGs). The proportions of positive selection, neutral evolution and purifying selection in the *A. halleri* lineage were compared in the two categories. In this test, the null model was the ratio of the two categories of OGGs (1,614 and 4,576 OGGs). The proportion of either positive selection or neutral evolution in genes duplicated only in the *A. halleri-lyrata* lineage was significantly higher than the proportion of either positive selection or neutral evolution in genes duplicated only in the *A. halleri* lineage (*P*-value = 5.1 × 10^-9^ for positive selection and 2.3 × 10^-5^ for neutral evolution by χ^2^ test, Fig. 4D). Conversely, the proportion of purifying selection in genes duplicated only in the *A. halleri-lyrata* lineage was significantly lower than the proportion of purifying selection in genes duplicated only in the *A. halleri* lineage (*P*-value = 2.2 × 10^-16^ by χ^2^ test, Fig. 4D). These results indicate that gene duplication in the *A. halleri-lyrata* lineage was the main determinant of the elevated proportion of positive selection in the *A. halleri* lineage.

### Functional bias of genes under positive or purifying selection in the A. halleri-lyrata and A. halleri lineages

We found that gene duplication contributed to functional divergence in comparison with non-duplicated genes in *Arabidopsis* but many duplicated genes had been retained with purifying selection, which induces an increase of gene dosage. Thus, we were interested in investigating the functional bias in duplicated and non-duplicated genes with positive/purifying selection in the *A. halleri-lyrata* and *A. halleri* lineages. OGGs were classified into duplicated and non-duplicated genes in the *A. halleri-lyrata* and *A. halleri* lineages. The OGGs were then further classified by positive and purifying selection. This gave a total of eight classes. In each class, we examined the degree of overrepresentation in 2,248 Gene Ontology (GO) categories compared with the number of GO categories assigned to *A. thaliana* genes (**see Materials and Methods**) **(Fig. 5A)**. Interestingly, the eight classes were largely clustered into two groups. The first group included four classes with positive selection. The second group included four classes with purifying selection. Each group was further classified by duplication and non-duplication regardless of lineage. Thus, genes with similar functional categories tended to have the same trend for gene duplication and/or selection pressure between the *A. halleri-lyrata* and *A. halleri* lineages, indicating that the evolutionary direction of *A. halleri* was quite similar in the *A. halleri-lyrata* and *A. halleri* lineages with respect to gene duplication and/or selection pressures.

**Figure 5.**
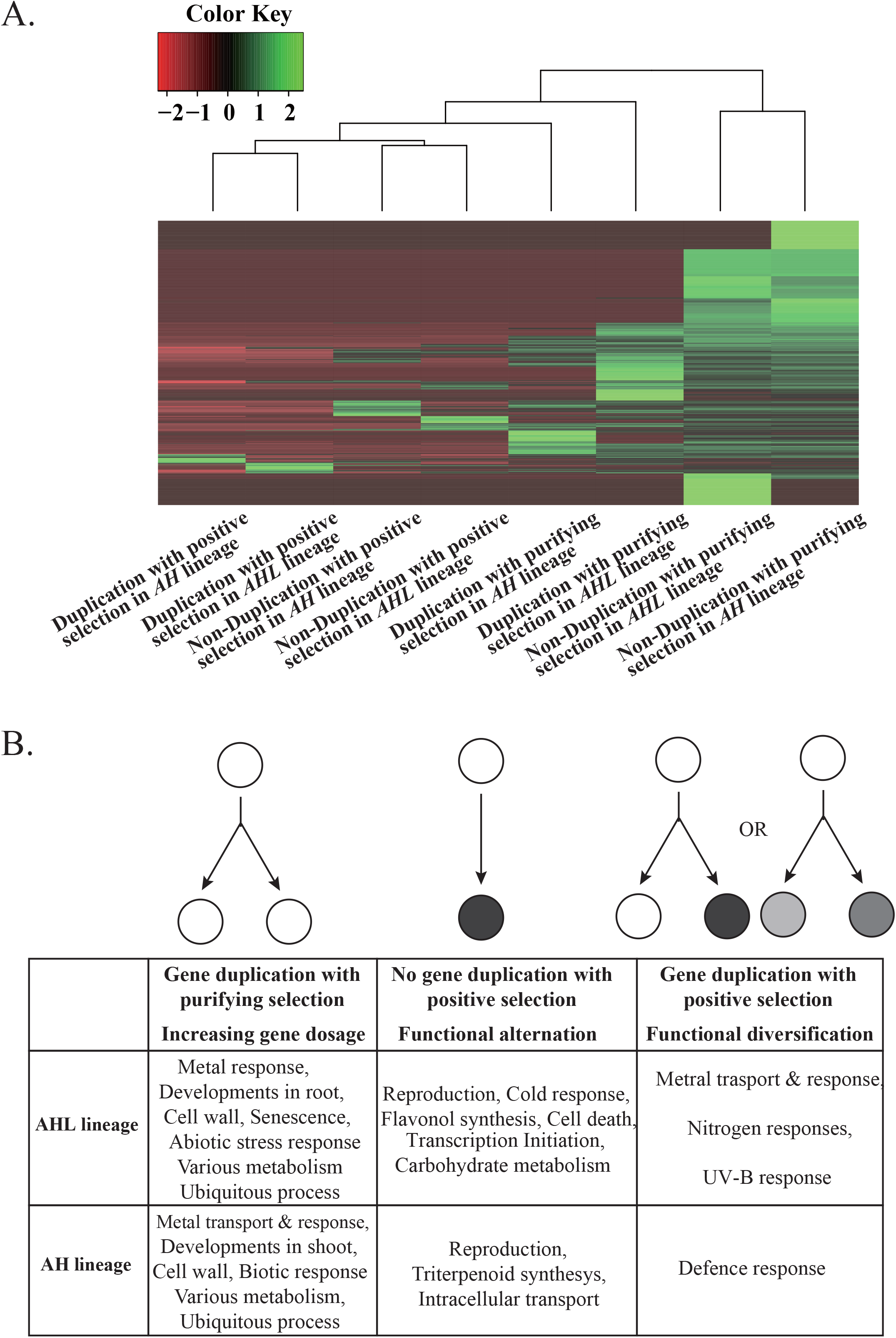
**Overrepresented functional categories** (A) The X-axis represents different kinds of OGGs. OGGs were classified into duplicated genes and non-duplicated genes in the *A. halleri-lyrata* (AHL) and *A. halleri* (AH) lineages. The OGGs were then further classified by positive or purifying selection. The Y-axis represents 2,248 Gene Ontology (GO) categories (biological processes) assigned to *A. thaliana* genes belonging to the OGGs. The key shows the relationship between colour and the z-score of overrepresentation in the GO categories. Red and green indicate low and high overrepresentation, respectively. The ratio of observed gene numbers in selected OGGs to expected gene numbers inferred from the data for all annotated genes was calculated in each GO category. (B) Overrepresented GO categories in OGGs with/without gene duplication and positive/purifying selection in the *A. halleri-lyrata* and *A. halleri* lineages (FDR < 0.01).

To examine the kinds of genes associated with phenotypic differences among *A. thaliana, A. lyrata* and *A. halleri*, we identified significantly overrepresented GO categories in OGGs with gene duplication and purifying selection, OGGs with gene duplication and positive selection and OGGs with non-duplication and positive selection in the *A. halleri-lyrata* and *A. halleri* lineages. OGGs with non-duplication and purifying selection were disregarded in the analysis because such genes tended to have the same functions in *A. halleri, A. lyrata* and *A. halleri*. From the GO categories assigned to the *A. thaliana* genes belonging to the OGGs, overrepresented GO categories were identified (**Figure 5B** and **Supplemental Table S2**; see Materials and Methods, FDR < 0.01).

When we focused on metal tolerance or accumulation among *A. thaliana*, *A. lyrata* and *A. halleri*, both *A. lyrata* and *A. halleri* had a higher tolerance to metal in comparison with *A. thaliana*. Metal tends to be accumulated in *A. halleri* compared with *A. lyrata*. This observation suggests that metal tolerance is enhanced in the *A. halleri-lyrata* lineage, and that metal accumulation is enhanced in the *A. halleri* lineage. Genes associated with metal responses and transporters were highly duplicated with purifying selection in the *A. halleri-lyrata* and *A. halleri* lineages, respectively, indicating that the dosages of genes associated with metal responses and transporters were enhanced in the *A. halleri-lyrata* and *A. halleri* lineages, respectively. Genes associated with metal responses were duplicated in the *A. halleri-lyrata* lineage and gene functions were altered without any duplication in the *A. halleri* lineage. Furthermore, genes associated with metal transport and responses were highly duplicated with positive selection in the *A. halleri-lyrata* lineage, indicating that functional divergence of genes associated with metal transport and responses occurred through gene duplication in the *A. halleri-lyrata* lineage. Thus, tolerance or accumulation of metal was enhanced in the *A. halleri-lyrata* and *A. halleri* lineages through gene duplication not only by enhancing gene dosage, but also by acquiring novel functions.

The other major phenotypic difference between *A. thaliana* and either *A. lyrata* or *A. halleri* is the reproductive system. *A. thaliana* has self-compatibility, but the others are self-incompatible. Genes associated with reproductive traits were subject to positive selection in the *A. halleri-lyrata* and *A. halleri* lineages without any gene duplication **(Fig. 5B, Supplemental Table S2)**, indicating that novel reproductive mechanisms tended to be lineage-specifically acquired through amino acid replacements only. Genes associated with UV and nitrogen responses tended to be duplicated with positive selection in the *A. halleri-lyrata* lineage **(Fig. 5B, Supplemental Table S2)**. Given that both *A. lyrata* and *A. halleri* can grow in harsher conditions, including sand dunes, cliffs, and high elevation sites, than *A. thaliana* (Mitchell-Olds 2001), *A. lyrata* and *A. halleri* may have more sophisticated mechanisms for surviving under low-nutrient soils or high UV. In the *A. halleri* lineage, genes associated with biotic responses tended to be duplicated with positive selection. It has been reported that lineage-specifically duplicated genes tend to be associated with biotic responses because pathogens tend to change rapidly in either evolutionary processes or environments. Thus, functional divergence of genes associated with biotic responses may have been advantageous during the evolution of *A. halleri* after its divergence from *A. lyrata*. Genes associated with cold responses, flavonol synthesis, cell death, transcription initiation, carbohydrate metabolism, triterpenoid synthesis and intracellular transport tended to have undergone positive selection without any gene duplication in both the *A. halleri-lyrata* and *A. halleri* lineages. However, we do not have any clear idea why these genes tended to show functional alteration.

Genes associated with development, the cell wall, senescence, abiotic responses, various metabolisms, biotic responses and ubiquitous processes tended to be duplicated in both the *A. halleri-lyrata* and *A. halleri* lineages with purifying selection **(Fig. 5B, Supplemental Table S2)**. These overrepresented functional categories may contribute to phenotypic differences between *A. halleri* and either *A. lyrata* or *A. thaliana* through high dosages. However, we do not know the phenotypic differences associated with these functional categories. These duplicated genes might have been retained because increased gene dosages associated with these functional categories were not too disadvantageous for *A. halleri*. To avoid gaining novel functions, these genes may be under purifying selection. In the future, these duplicated genes may be lost if disadvantageous functions appear. Indeed, the *A. halleri-lyrata* and *A. halleri* lineages have extraordinarily high retention rates of duplicated genes in comparison with earlier plant lineages. These observations indicate that most of the duplicated genes in the *A. halleri-lyrata* and *A. halleri* lineages may be lost in future evolution.

### Concluding remarks

In the present analysis, we generated 28,036 *A. halleri* genes that were orthologous to 86% of the annotated *A. lyrata* genes from contigs generated from Illumina short reads. From 28,036 OGGs between *A. lyrata* and *A. halleri*, we generated 23,086 OGGs among *A. halleri, A. lyrata* and *A. thaliana*, and identified 3,268 gain events from gene duplication in the *A. halleri-lyrata* lineage. Furthermore, using the mapping coverage of the Illumina reads, we identified 3,741 gain events from gene duplication in the *A. halleri* lineage. The gain rates were inferred to be 2.0–2.3 and 5.7–6.7 × 10^-2^ per gene per MY in the *A. halleri-lyrata* and *A. halleri* lineages, respectively. The gain rate in the *A. halleri* lineage was approximately two to three times higher than in the *A. halleri-lyrata* lineage. Although the gain events in *A. halleri* were inferred from Illumina reads, the inferred gain rate is reasonable because the number of gain events in the *A. lyrata* lineage after it split from *A. halleri* was 3,512, which is equivalent to the number of gene gains in the *A. halleri* lineage. Using our previous data, we re-estimated the gain rates in the three time periods after the divergence of mosses, rice and poplar (**Fig. 1**). The inferred gain rates (1.8–3.0 × 10^-3^) were approximately 10–30 times lower than in the *Arabidopsis* lineage (**Fig. 1**). Thus, gain rates tend to increase as the evolutionary period gets younger. One explanation for this gain rate difference is that duplication rates might be substantially higher immediately after speciation. Another explanation is that duplicated genes tend to rapidly decay over time. This explanation is supported by several previous reports in which younger duplicated genes tended to be relaxed compared with older duplicated genes (Lynch and Conery 2000; Jordan et al. 2004; Alba and Castresana 2005; Wolf et al. 2009; Vishnoi et al. 2010). That is, most anciently duplicated genes tend not to be retained in current species. Consequently, the gain rates inferred in earlier evolutional periods tend to decrease.

To investigate the functional divergence of duplicated genes in the *A. halleri-lyrata* and *A. halleri* lineages, we identified OGGs under either positive or purifying selection in these lineages based on the ratio of nonsynonymous and synonymous substitution rates (*K*_A_/*K*_S_). Interestingly, the proportions of positive and purifying selection tended to increase and decrease, respectively, when gene duplication occurred in either the *A. halleri-lyrata* or *A. halleri* lineage. This result indicates that gene duplication tends to enhance functional divergence in comparison with non-duplicated genes in the *Arabidopsis* lineage. In contrast, the general observation is that duplicated genes tend to have less functional divergence in yeasts, plants and mammals (Yang et al. 2003; Davis and Petrov 2004; Yang and Gaut 2011). This is because functionally important genes are more likely to be retained as duplicates (Davis and Petrov 2004). This contradictory relationship might derive from the duplication ages. Most previous analyses have examined recently observed selective pressures in anciently duplicated genes. When functional divergence was examined in recently duplicated genes, the duplicated genes tended to have higher functional divergence than singletons (Satake et al. 2012). Together, these results suggest duplicated genes tend to have higher functional divergence immediately after duplication than singletons.

How long gene duplication accelerates functional divergence remains an open question. To address this, we examined whether gene duplication in the *A. halleri-lyrata* lineage (2–10 MYA) accelerated functional divergence in the *A. halleri* lineage (<2 MYA). Interestingly, we found that gene duplication in the *A. halleri-lyrata* lineage enhanced the proportion of positive selection in the *A. halleri* lineage with a rate approximately 2.5 times higher than in the *A. halleri* lineage **(Fig. 4C and 4D)**. This result indicated that the functional divergence of duplicated genes was accelerated several MY after gene duplication. If gene duplication is too deleterious for a gene, the gene tends to be lost immediately after duplication. If not, duplicated genes may be retained for a long period without functional divergence because functional divergence may be evolutionarily disadvantageous. Therefore, immediately after duplication, most duplicated genes might be under functional constraints. Indeed, many recently duplicated genes have functional redundancy in *A. thaliana* (Hanada et al. 2009a; Hanada et al. 2011). These young duplicated genes tend to be less functionally constrained than singletons, and may have the potential to obtain an essential function to survive in new environments.

Finally, we examined the kinds of genes that were duplicated and/or under positive selection in the *A. halleri-lyrata* and *A. halleri* lineages. Although similar functional categories tended to have experienced gene duplication and/or selection pressure in the *A. halleri-lyrata* and *A. halleri* lineages, we found that genes associated with differential traits among *A. halleri*, *A. lyrata* and *A. thaliana* tended to have undergone gene duplication and/or positive selection in the *A. halleri-lyrata* and *A. halleri* lineages. For example, *A. halleri* is known as a heavy metal hyper-accumulator with high metal tolerance. *A. lyrata* is tolerant of heavy metal iron in the soil to some degree but *A. thaliana* is not. Genes related to heavy metal tolerance and accumulation tended to be highly duplicated with positive or purifying selection in the *A. halleri-lyrata* and *A. halleri* lineages. Earlier studies reported that metal tolerance was enhanced by increasing gene dosage through gene duplication (Hanikenne et al. 2008). However, it is likely that metal tolerance and accumulation were acquired through novel functions via gene duplications in *A. halleri*. Taken together, the results of our study reveal that lineage-specifically duplicated genes have contributed to species-specific evolution in *Arabidopsis*.

## Methods

### Sampling and Illumina sequencing analysis

*Arabidopsis halleri* subsp. *gemmifera* is one of the three subspecies of *A. halleri* that grows in the Russian Far East, northeastern China, Korea, Taiwan, and Japan (Al-Shehbaz and O’Kane Jr 2002). In 2008, a leaf sample was collected from an individual of *A. halleri* subsp. *gemmifera* on Mt. Ibuki, Japan. DNA was extracted from the leaf using a DNeasy Plant Mini Kit (QIAGEN, Venlo, Netherlands). A 300-bp paired-end DNA library was constructed according to the Paired-End Genomic DNA Sample Preparation Kit (Illumina, San Diego, California, U.S.), and 93-bp paired-end reads were obtained from the Illumina GAIIx sequencer.

### Assembly of A. halleri genes orthologous to A. lyrata genes

A total of 44.5 Gb reads were determined by Illumina DNA paired-end sequencing. Approximately 28% of reads had either low-quality scores or adapters and were trimmed by Trim Galore (www.bioinformatics.babraham.ac.uk). When a paired-end read was completely removed by this procedure, the other read was used as a single-end read. Given that mitochondrial and chloroplast genomes have much higher copy numbers than the nuclear genome, sequencing reads mapped to either the mitochondrial or chloroplast genome in *A. thaliana* (https://www.arabidopsis.org, TAIR10) or *A. lyrata* (http://genome.jgi.doe.gov, FilteredModels6) by BLASTN were excluded from the following procedures (Boratyn et al. 2013). Assuming that the genome size of *A. halleri* was 255 Mb (Johnston et al. 2005), the sequencing coverage was estimated to be approximately 135× (34.4/0.255 Gb). The sequencing reads were assembled with either the ABySS software (Simpson et al. 2009), SOAP-denovo2 (Luo et al. 2012) or CLC Genomics Workbench 7.0.3 (http://www.clcbio.com). Genes in close species tend to be mapped to reliable assembled sequences with a higher proportion. As the closest species, 32,670 annotated *A. lyrata* genes (http://genome.jgi.doe.gov, FilteredModels6) were mapped to each type of assembled DNA segment by the gmap software, which takes into account exon-intron junctions (Wu and Watanabe 2005). The number of mapped *A. lyrata* genes (>80% coverage) was 23,681 for the assembled DNA segments generated by ABySS with 63 *K*-mer size, which was the highest proportion among the different types of assembled DNA segments. When a mapped sequence had more than 80% similarity and 80% coverage against an *A. halleri* gene, it was defined as an orthologous gene of *A. halleri* against *A. lyrata*. To collect additional *A. halleri* genes orthologous to *A. lyrata*, unmapped *A. lyrata* genes were re-mapped to the assembled DNA segments by TBLASTN (Boratyn et al. 2013). When an *A. lyrata* gene was mapped to more than two contigs, the contigs were concatenated following the direction of the *A. lyrata* gene. The *A. lyrata* gene was then mapped to the concatenated contigs. We found 4,355 mapped sequences with more than 80% similarity and 80% coverage against *A. halleri* genes. Finally, we succeeded in identifying 28,036 (85.8%) *A. halleri* genes orthologous to 32,670 *A. lyrata* genes. The analysis procedures and findings are summarized in **Supplemental Figure S1**.

### Construction of orthologous gene groups among B. rapa, A. thaliana, A. lyrata and A. halleri genes

There were 28,036 pairs of orthologous genes between *A. halleri* and *A. lyrata*. A similarity search of the translated sequences of the *A. halleri* and *A. lyrata* genes was performed for annotated representative *A. thaliana* genes (TAIR10) with BLASTP (Boratyn et al. 2013). When both the *A. halleri* and *A. lyrata* genes in an orthologous pair had hits to the same *A. thaliana* gene with the best score, the three genes were categorized into the same orthologous group. Thus, we identified 24,784 orthologous groups including *A. thaliana, A. lyrata* and *A. halleri* genes. Among these, we searched for orthologous groups that were consistent with the species tree. To do this, the synonymous substitution rates (*K*_S_) were calculated among the *A. thaliana, A. lyrata* and *A. halleri* genes in each orthologous group by yn00 in PAML (Yang 2007). When the *K*_S_ between *A. lyrata* and *A. halleri* was lower than either the *K*_S_ between *A. thaliana* and *A. lyrata* or the *K*_S_ between *A. thaliana* and *A. halleri*, we assumed that the topology of the orthologous group was consistent with the species tree. Out of 24,784 orthologous groups, we identified 23,086 (23,086/24,956 = 92.5%) orthologous groups that followed the speciation process among *A. thaliana, A. lyrata* and *A. halleri* genes. The analysis procedures and findings are summarized in **Supplemental Figure S2**.

To identify orthologous *Brassica rapa* genes to the 23,086 orthologous groups, we downloaded the *B. rapa* genes from Phytozome (version 10: BrapaFPsc_277_v1.3) (Goodstein et al. 2012). Similarity searches of the translated sequences of the *A. thaliana, A. halleri* and *A. lyrata* genes were performed for the translated sequences of the *B. rapa* genes with BLASTP (Boratyn et al. 2013). When the *A. thaliana, A. lyrata* and *A. halleri* genes in an orthologous pair had hits to the same *B. rapa* gene with the best score, the four genes were categorized into the same orthologous group. Thus, we identified 20,366 orthologous groups including *B. rapa, A. thaliana, A. lyrata* and *A. halleri* genes. Among these, we searched for orthologous groups that were consistent with the species tree. To do this, we generated a phylogenetic tree by the neighbor-joining method using the PAUP software (set outroot = mono, dset distance = hky) (Saitou and Nei 1987; Wilgenbusch and Swofford 2003). When the topology of the gene tree was the same as that of the species tree, we assumed that the topology of the orthologous group was consistent with the species tree. Out of 20,366 orthologous groups, we identified 17,604 orthologous groups that followed the speciation process among *B. rapa, A. thaliana, A. lyrata* and *A. halleri* genes. The analysis procedures and findings are summarized in **Supplemental Figure S3**.

### Validation of A. halleri lineage-specifically duplicated genes by droplet digital PCR

The Illumina sequencing reads were mapped to 28,036 *A. halleri* genes by BOWTIE2 (Langdon 2015). To calculate the read coverage of each gene, the mapped count was divided by the number of genes to which a read was mapped. Reads per kilobase of exon model per million (RPKM) values were calculated for each *A. halleri* gene. For 11 *A. halleri* genes whose copy numbers were known (**Table S5**), we designed primer pairs using the following parameters in the Primer3Plus software (Untergasser et al. 2007): primer length of 18–24 bases, amplicon length of 70–150 bp, *T*_m_ value of 57–63°C and GC composition of 40–60% (**Table S6**). To obtain enough genomic DNA for ddPCR, we mixed the genomic DNAs from four individuals of *A. halleri* from Mt. Ibuki. The genomic DNA was sonicated with the Covaris M220 system (25 s in frequency sweeping mode). The concentration of the sonicated genomic DNA sample was 6 ng/μl. The peak size of sonicated DNA fragments was 2382 bp according to the Fragment Analyzer™ system (Advanced analytical, Ankeny, USA) with the High Sensitivity Genomic DNA Analysis Kit (Advanced analytical). Each ddPCR reaction was performed with ddPCR EvaGreen Supermix (Bio-Rad, California, USA). Each reagent was divided into approximately 20,000 droplets with a droplet generator (Bio-Rad QX-200). Cycled droplets were measured with a QX200 droplet reader (Bio-Rad).

To find *A. halleri* genes whose DNA concentrations were robustly inferred by ddPCR, we first identified uniquely mapped regions (>80 bp) in *A. halleri* genes from the Illumina sequencing reads. Among the *A. halleri* genes with uniquely mapped regions, we manually chose 50 *A. halleri* genes with a variety of RPKMs. We designed a pair of primers for each gene according to the following parameters in the Primer3Plus software (Untergasser et al. 2007): primer length of 18–24 bases, amplicon length of 70–150 bp, *T*_m_ value of 57–63°C and GC composition of 40–60%. To examine the specificity of the targeted DNAs, we performed real-time PCR analysis using the protocol for the Mx3000P qPCR System (Agilent Technologies). The PCR analyses were performed using SsoFast EvaGreen Supermix (Bio-Rad) and the products were analyzed using the Mx3000P multiplex quantitative PCR system (Agilent Technologies). Specific and multiple reactions should result in a single and multiple melting peaks corresponding to the PCR product. Of the 50 primer pairs, 27 showed a clear raised curve for all three replicates. Thus, for the copy number assay by ddPCR, we used 27 primer pairs for unknown copy number genes, 11 pairs for single-copy genes in a broad range of plant species and 2 pairs for a three-copy gene.

### Selective pressures in the A. halleri-lyrata and A. halleri lineages

To infer selective pressure in the *A. halleri-lyrata* lineage, we focused on 17,641 orthologous groups that followed the speciation process among *B. rapa, A. thaliana, A. lyrata* and *A. halleri* genes. In each orthologous group, a multiple alignment was made to match the coding regions with the computer program MAFFT (Katoh et al. 2002). Using the phylogenetic tree and multiple alignment, we constructed the common ancestral sequences among *A. thaliana*, *A. halleri* and *A. lyrata*, and the common ancestral sequence between *A. halleri* and *A. lyrata* by codeml (model = 1, NSsites = 0) in PAML. For each pair of ancestral sequences, the synonymous (*K*_S_) and nonsynonymous substitution rates (*K*_A_) were calculated by yn00 (icode = 0, weighting = 0, commonf3×4 = 0) in PAML. To determine whether the *K*_A_/*K*_S_ ratio was significantly <1, two maximum likelihood values were calculated with the *K_A_/K_S_* ratio fixed at 1 and with the *K_A_/K_S_* ratio as a free parameter. The ratio of the maximum likelihood values was then compared to the χ^2^ distribution. A *P*-value representing the deviation of the two models was then calculated for the *A. halleri-lyrata* lineage in each of the 17,641 orthologous groups.

To infer selective pressure in the *A. halleri* lineage, we generated a tree file trifurcating among the *A. thaliana*, *A. lyrata* and *A. halleri* genes in 23,086 OGGs. In each of the orthologous groups, a multiple alignment was made to match the coding regions by the computer program MAFFT (Katoh et al. 2002). Using the tree file and multiple alignment file, we constructed the common ancestral sequences among *A. thaliana*, *A. halleri* and *A. lyrata* by codeml (model = 1, NSsites = 0) in PAML. Although we used a representative *A. halleri* gene sequence to infer the ancestral sequence, proper *A. halleri* genes had sequence variation when they were lineage-specifically duplicated after the split from *A. lyrata*. From the Illumina sequencing reads mapped to 28,036 *A. halleri* genes by BOWTIE2 (Langdon 2015), the sequence variation was examined in each *A. halleri* gene by the Picard software (http://broadinstitute.github.io/picard/). When a variable sequence was observed in the *A. halleri* genes, the codon sequence including the variable nucleotide was concatenated into the representative *A. halleri* genes. Alignments between *A. halleri* genes including codons with variable nucleotides and the ancestral sequence were modified by adding codons of the ancestral sequence against concatenated codons including variable nucleotides. *K_A_* and *K_S_* were calculated in each pair of ancestral and *A. halleri* gene sequences including variable sequences by yn00 in PAML. To determine whether the *K_A_/K_S_* ratio was significantly <1, two maximum likelihood values were calculated with the *K_A_/K_S_* ratio fixed at 1 and with the *K_A_/K_S_* ratio as a free parameter by codeml in PAML. The ratio of the maximum likelihood values was then compared to the χ^2^ distribution. A *P*-value representing the deviation of the two models was then calculated for each *A. halleri* gene. The analysis procedures are summarized in **Supplemental Figure S4**.

### Gene annotation of A. halleri genes

GO assignments for the *Arabidopsis* genes were obtained from The *Arabidopsis* Information Resource (www.arabidopsis.org). Among the top three GO categories (cellular components, molecular functions, and biological processes), we analyzed only biological processes. For the GO categories belonging to the biological processes category, we recorded the numbers of *Arabidopsis* genes that were assigned and not assigned to OGGs. In each GO category, the expected values were compared with the observed values using a χ^2^ test to determine whether the ratio of observed gene numbers in the assigned genes to those in the non-assigned genes was significantly higher than the expected ratio. The ratio of observed gene numbers to expected gene numbers was calculated in each GO category for different categories of OGGs. The ratios were processed in the R software environment (www.r-project.org). The ratios were normalized among different arrays using Z scores calculated by R library genescale. HeatMap was generated by R library heatmap.2. To correct for multiple testing, the FDR was estimated by the R-library Q-VALUE software. The null hypothesis was rejected if FDR values were <0.01.

## Data Access

Next-generation sequence data generated for this study have been submitted to D-way (https://trace.ddbj.nig.ac.jp/D-way/) under the accession number DRA004564.

## Acknowledgments

We thank Kiyomi Imamura, Makiko Tosaka, Taiji Kikuchi, Terumi Horiuchi, Kanako Onizuka and Miu Kubota for Illumina sequencing analyses and ddPCR analyses. We also thank the National Institute of Genetics of the Research Organization of Information and Systems for providing excellent supercomputer services. This work was supported by the Research for Evolutional Science and Technology (CREST) program “Creation of essential technologies to utilize carbon dioxide as a resource through the enhancement of plant productivity and the exploitation of plant products” of the Japan Science and Technology Agency (JST) (K.H. and S.I.M.); Grants-in-Aid for Scientific Research (to K.H. and S.I.M.).

## Author contributions

KH and SIM designed the studies. KH performed data analyses and wrote the manuscript. AT and MN performed ddPCR analysis and helped to write the manuscript. YS and SS performed Illumina sequencing analysis. AJN and MI helped to sample *Arabidopsis halleri*. SIM provided experimental input and helped to write the manuscript.

## REFERENCE

Al-Shehbaz IA, O'Kane Jr SL. 2002. Taxonomy and phylogeny of Arabidopsis (Brassicaceae). The Arabidopsis book/American Society of Plant Biologists 1.

Alba MM, Castresana J. 2005. Inverse relationship between evolutionary rate and age of mammalian genes. Mol Biol Evol 22: 598–606.

Arabidopsis_Genome_Initiative. 2000. Analysis of the genome sequence of the flowering plant Arabidopsis thaliana. Nature 408: 796–815.

Beilstein MA, Nagalingum NS, Clements MD, Manchester SR, Mathews S. 2010. Dated molecular phylogenies indicate a Miocene origin for Arabidopsis thaliana. Proc Natl Acad Sci U S A 107: 18724–18728.

Boratyn GM, Camacho C, Cooper PS, Coulouris G, Fong A, Ma N, Madden TL, Matten WT, McGinnis SD, Merezhuk Y et al. 2013. BLAST: a more efficient report with usability improvements. Nucleic acids research 41: W29–33.

Clark RM, Schweikert G, Toomajian C, Ossowski S, Zeller G, Shinn P, Warthmann N, Hu TT, Fu G, Hinds DA et al. 2007. Common sequence polymorphisms shaping genetic diversity in Arabidopsis thaliana. Science 317: 338–342.

Davis JC, Petrov DA. 2004. Preferential duplication of conserved proteins in eukaryotic genomes. PLoS Biol 2: E55.

Duarte JM, Wall PK, Edger PP, Landherr LL, Ma H, Pires JC, Leebens-Mack J, dePamphilis CW. 2010. Identification of shared single copy nuclear genes in Arabidopsis, Populus, Vitis and Oryza and their phylogenetic utility across various taxonomic levels. BMC evolutionary biology 10: 61.

Fortna A, Kim Y, MacLaren E, Marshall K, Hahn G, Meltesen L, Brenton M, Hink R, Burgers S, Hernandez-Boussard T et al. 2004. Lineage-specific gene duplication and loss in human and great ape evolution. PLoS Biol 2: E207.

Goodstein DM, Shu S, Howson R, Neupane R, Hayes RD, Fazo J, Mitros T, Dirks W, Hellsten U, Putnam N et al. 2012. Phytozome: a comparative platform for green plant genomics. Nucleic acids research 40: D1178–1186.

Hanada K, Kuromori T, Myouga F, Toyoda T, Li WH, Shinozaki K. 2009a. Evolutionary persistence of functional compensation by duplicate genes in Arabidopsis. Genome Biol Evol 1: 409–414.

Hanada K, Kuromori T, Myouga F, Toyoda T, Shinozaki K. 2009b. Increased expression and protein divergence in duplicate genes is associated with morphological diversification. PLoS Genet 5: e1000781.

Hanada K, Sawada Y, Kuromori T, Klausnitzer R, Saito K, Toyoda T, Shinozaki K, Li WH, Hirai MY. 2011. Functional compensation of primary and secondary metabolites by duplicate genes in Arabidopsis thaliana. Mol Biol Evol 28: 377–382.

Hanada K, Vallejo V, Nobuta K, Slotkin RK, Lisch D, Meyers BC, Shiu SH, Jiang N. 2009c. The functional role of pack-MULEs in rice inferred from purifying selection and expression profile. Plant Cell 21: 25–38.

Hanada K, Zou C, Lehti-Shiu MD, Shinozaki K, Shiu SH. 2008. Importance of lineage-specific expansion of plant tandem duplicates in the adaptive response to environmental stimuli. Plant Physiol 148: 993–1003.

Hanikenne M, Talke IN, Haydon MJ, Lanz C, Nolte A, Motte P, Kroymann J, Weigel D, Kramer U. 2008. Evolution of metal hyperaccumulation required cis-regulatory changes and triplication of HMA4. Nature 453: 391–395.

Heckman TM, Kauffmann G. 2011. The coevolution of galaxies and supermassive black holes: a local perspective. Science 333: 182–185.

Heredia NJ, Belgrader P, Wang S, Koehler R, Regan J, Cosman AM, Saxonov S, Hindson B, Tanner SC, Brown AS et al. 2013. Droplet Digital PCR quantitation of HER2 expression in FFPE breast cancer samples. Methods 59: S20–23.

Hu TT, Pattyn P, Bakker EG, Cao J, Cheng JF, Clark RM, Fahlgren N, Fawcett JA, Grimwood J, Gundlach H et al. 2011. The Arabidopsis lyrata genome sequence and the basis of rapid genome size change. Nature genetics 43: 476–481.

Johnston JS, Pepper AE, Hall AE, Chen ZJ, Hodnett G, Drabek J, Lopez R, Price HJ. 2005. Evolution of genome size in Brassicaceae. Ann Bot 95: 229–235.

Jordan IK, Wolf YI, Koonin EV. 2004. Duplicated genes evolve slower than singletons despite the initial rate increase. BMC evolutionary biology 4: 22.

Katoh K, Misawa K, Kuma K, Miyata T. 2002. MAFFT: a novel method for rapid multiple sequence alignment based on fast Fourier transform. Nucleic acids research 30: 3059–3066.

Koenig D, Weigel D. 2015. Beyond the thale: comparative genomics and genetics of Arabidopsis relatives. Nat Rev Genet 16: 285–298.

Kramer U. 2010. Metal hyperaccumulation in plants. Annual review of plant biology 61: 517–534.

Langdon WB. 2015. Performance of genetic programming optimised Bowtie2 on genome comparison and analytic testing (GCAT) benchmarks. BioData Min 8: 1.

Lockton S, Gaut BS. 2005. Plant conserved non-coding sequences and paralogue evolution. Trends Genet 21: 60–65.

Luo R, Liu B, Xie Y, Li Z, Huang W, Yuan J, He G, Chen Y, Pan Q, Liu Y et al. 2012. SOAPdenovo2: an empirically improved memory-efficient short-read de novo assembler. Gigascience 1: 18.

Lynch M, Conery JS. 2000. The evolutionary fate and consequences of duplicate genes. Science 290: 1151–1155.

Mitchell-Olds T. 2001. Arabidopsis thaliana and its wild relatives: a model system for ecology and evolution. Trends in Ecology and Ecolution 16: 693–700.

Moghe GD, Hufnagel DE, Tang H, Xiao Y, Dworkin I, Town CD, Conner JK, Shiu SH. 2014. Consequences of Whole-Genome Triplication as Revealed by Comparative Genomic Analyses of the Wild Radish Raphanus raphanistrum and Three Other Brassicaceae Species. Plant Cell 26: 1925–1937.

Rensing SA, Lang D, Zimmer AD, Terry A, Salamov A, Shapiro H, Nishiyama T, Perroud PF, Lindquist EA, Kamisugi Y et al. 2008. The Physcomitrella genome reveals evolutionary insights into the conquest of land by plants. Science 319: 64–69.

Rizzon C, Ponger L, Gaut BS. 2006. Striking similarities in the genomic distribution of tandemly arrayed genes in Arabidopsis and rice. PLoS Comput Biol 2: e115.

Rostoks N, Borevitz JO, Hedley PE, Russell J, Mudie S, Morris J, Cardle L, Marshall DF, Waugh R. 2005. Single-feature polymorphism discovery in the barley transcriptome. Genome Biol 6: R54.

Saitou N, Nei M. 1987. The neighbor-joining method: a new method for reconstructing phylogenetic trees. Mol Biol Evol 4: 406–425.

Satake M, Kawata M, McLysaght A, Makino T. 2012. Evolution of vertebrate tissues driven by differential modes of gene duplication. DNA Res 19: 305–316.

Shimizu KK, Purugganan MD. 2005. Evolutionary and ecological genomics of Arabidopsis. Plant Physiol 138: 578–584.

Simpson JT, Wong K, Jackman SD, Schein JE, Jones SJ, Birol I. 2009. ABySS: a parallel assembler for short read sequence data. Genome research 19: 1117–1123.

Turner TL, Bourne EC, Von Wettberg EJ, Hu TT, Nuzhdin SV. 2010. Population resequencing reveals local adaptation of Arabidopsis lyrata to serpentine soils. Nat Genet 42: 260–263.

Tuskan GA Difazio S Jansson S Bohlmann J Grigoriev I Hellsten U Putnam N Ralph S Rombauts S Salamov A et al. 2006. The genome of black cottonwood, Populus trichocarpa (Torr. & Gray). Science 313: 1596–1604.

Untergasser A, Nijveen H, Rao X, Bisseling T, Geurts R, Leunissen JA. 2007. Primer3Plus, an enhanced web interface to Primer3. Nucleic Acids Res 35: W71–74.

Vanneste K, Baele G, Maere S, Van de Peer Y. 2014. Analysis of 41 plant genomes supports a wave of successful genome duplications in association with the Cretaceous-Paleogene boundary. Genome research 24: 1334–1347.

Verbruggen N, Hermans C, Schat H. 2009. Molecular mechanisms of metal hyperaccumulation in plants. New Phytol 181: 759–776.

Vishnoi A, Kryazhimskiy S, Bazykin GA, Hannenhalli S, Plotkin JB. 2010. Young proteins experience more variable selection pressures than old proteins. Genome research 20: 1574–1581.

Wang J, Marowsky NC, Fan C. 2013. Divergent evolutionary and expression patterns between lineage specific new duplicate genes and their parental paralogs in Arabidopsis thaliana. PLoS One 8: e72362.

Wilgenbusch JC, Swofford D. 2003. Inferring evolutionary trees with PAUP*. Curr Protoc Bioinformatics **Chapter 6**: Unit 6 4.

Wolf YI, Novichkov PS, Karev GP, Koonin EV, Lipman DJ. 2009. The universal distribution of evolutionary rates of genes and distinct characteristics of eukaryotic genes of different apparent ages. Proc Natl Acad Sci U S A 106: 7273–7280.

Wolfe KH, Gouy M, Yang YW, Sharp PM, Li WH. 1989. Date of the monocot-dicot divergence estimated from chloroplast DNA sequence data. Proc Natl Acad Sci U S A 86: 6201–6205.

Wu TD, Watanabe CK. 2005. GMAP: a genomic mapping and alignment program for mRNA and EST sequences. Bioinformatics 21: 1859–1875.

Yang J, Gu Z, Li WH. 2003. Rate of protein evolution versus fitness effect of gene deletion. Mol Biol Evol 20: 772–774.

Yang L, Gaut BS. 2011. Factors that contribute to variation in evolutionary rate among Arabidopsis genes. Mol Biol Evol 28: 2359–2369.

Yang Z. 2007. PAML 4: phylogenetic analysis by maximum likelihood. Mol Biol Evol 24: 1586–1591.

